# Following the evolution of *Homo sapiens* across Africa using a uniparental genetic guide

**DOI:** 10.1101/2022.07.06.499026

**Authors:** Vicente M. Cabrera

## Abstract

The origin and evolution of modern humans in Africa has reached a multidisciplinary consensus but the age and regions where it originated and evolved are current topics of discussion. In this study I put forward an integrative model guided by the phylogeny and phylogeography of mitochondrial DNA (and Y-chromosome) haplogroups. I propose an early origin of modern humans in northwest Africa in a temporal window of 257-345 thousand years ago. A first population split in central Africa around 175-288 thousand years ago. A subsequent northward spread with additional population subdivisions during a long statistical interval that culminated in a first successful out of Africa migration around 130 thousand years ago. A population constriction in southwest Asia motivated an early return to Africa between 79 and 107 thousand years ago. This ample Eurasian-ebb to Africa, detected by mitochondrial haplogroup L3 and Y-chromosome haplogroup E preceded other later and geographically more limited Eurasian backflows. The archaeological and fossil finds that could be coetaneous to this molecular journey have been integrated into this interdisciplinary model.

## Introduction

Hypotheses about human evolution, formulated from archaeological and genetics data, are mainly based on radiometric dating for the former and on molecular dating for the latter [1]. These methods have the advantage of locating important evolutionary events in specific places and time frames where these events must have occurred. However, in many cases, the frameworks established by different disciplines conflict. For example, applying the molecular clock to mitochondrial DNA (mtDNA) genetic variation, it has been established that modern humans had a genetic African origin around 200 thousand years ago (kya) [2], and that a more evolved form of that lineage left Africa colonizing Eurasia around 60 kya [3]. However, Middle Stone Age (MSA) artefacts and fossils dated at the site of Jebel Irhoud, Morocco placed the *Homo sapiens* emergence in northwest Africa around 300 kya [4,5], and roughly at the same time MSA tool assemblages replaced more primitive Acheulean tools in southern Kenya [6,7]. Furthermore, fossils from Misliya Cave, Israel, dated around 180 kya [8], and in southern China dated around 100 kya [9], strongly suggest that members of the *H. sapiens* clade left Africa earlier than previously thought. It could be adduced that because mtDNA is a single inherited female locus, its chronology might be discordant with those obtained from whole nuclear genome analysis. However, with few exceptions [10], the human demographic history deduced from genomic studies highly resembles the one based on uniparental markers [11,12]. Therefore, the most prevalent opinion from the population genetics field is that demographic human expansions from Africa to the Middle East and beyond, prior to approximately 60 Kya, were ephemeral dispersals that did not contributed to the modern human gene pool [13]. However, the genetic chronological framework is based on an insecure evolutionary rate, which in turn depends on the germline mutation rate, selective constrains, and the fluctuation of the effective population size due to demographic processes [14]. Certainly, new technological progresses in DNA sequencing have highly refined the human germline mutation rate both at the whole genome [15] and the mtDNA levels [16,17]. In addition, purifying selection has been taken into account to improve evolutionary rate estimations [18], but it seems that a time-dependence effect on this rate [19,20], most probably due to fluctuations in population size [14], is the main factor responsible of the changes detected in the evolutionary rate values observed. Recently, a simple algorithm has been proposed to counterbalance these effects on the mtDNA genome, which practically doubles the coalescent time estimations along the human mtDNA phylogenetic tree [21]. In this way, the most determinant archaeological findings related to the human evolution coherently fit into the molecular chronology [21].

In this paper, using that algorithm, with an appropriate germline mutation rate, and the successive coalescent events across the human mtDNA phylogeny as a molecular guide, I describe the progressive evolution of modern humans into Africa using an integrative model that incorporates the main archaeological and genetic evolutionary discoveries into a coherent picture.

## Material and methods

### Material

For the phylogenetic and phylogeographic analyses I searched for mtDNA complete genomes at the NCBI GenBank (www.ncbi.nlm.nih.gov/genbank/), and MITOMAP (www.mitomap.org/MITOMAP) databases, choosing representatives of all African haplogroups and their main subgroups. Sequences were classified according to the PhyloTree v.17 (http://www.phylotree.org) [22]. In total I analysed 1,010 mitogenomes (243 for L0, 140 for L1, 73 for L5, 210 for L2, 8 for L6, 32 for L4, and 304 for L3). GenBank accession numbers for these sequences, their haplogroup classification, and their country/ethnic affiliation are detailed in supplementary (S) Table 1. A phylogenetic tree showing the major mtDNA haplogroups relationships is presented as supplementary(S) Fig 1. Phylogeographic trees for haplogroups L2, L6 and L4 are presented in SFig 2, 3, and 4 respectively. The phylogeography of haplogroups L0, L1, L5, and L3 have been studied previously [23].

### Methods

Phylogenetic trees were built using median-joining networks [24]. To calculate coalescent absolute ages of the main haplogroups I used a mutation rate of 1.6 x 10^-8^ per site per year (assuming a mtDNA genomic length of 16,500 base pairs) that is the mean of two independent empirical estimates [16,17], and applied a composite rho algorithm that takes into account time-dependence effects on this mutation rate [21]. The procedure for obtaining coalescent ages for the main haplogroups L0, L1, L5, L2, L6, L4, and L3 are detailed in STables 2, 3, 4, 5, 6, 7, and 8 respectively. For relative age comparisons of phylogeographically representative subclades, I calculated their coalescent age using the rho statistic [25] and a mutation rate for the complete mtDNA sequence of one substitution in every 3,624 years, correcting for purifying selection [18], but using those sequences with the largest number of mutations within each clade. The reason of this is that the effects of both selection (mainly purifying selection) and genetic drift tend to eliminate those sequences that in a Poisson distribution, with very low mean of success, have a greater number of mutations and conserve those included in the largest classes that have zero or very few mutations [14].

## Results

### The genus *Homo* from a genetic perspective

*Homo* is a genus represented by only a single extant species (modern humans) and several extinct specimens [26], which appeared during an interval of just over two million years. The first hominin species with worldwide spread, most probably as result of consecutive waves of expansion [27], was *Homo erectus s.l.* Remains of this species have been unearthed in Africa [28], the Middle East [29], the Caucasus [30], China [31] and Indonesia [32]. As a generalist species, *H. erectus* reached this wide geographic range with migrant groups adapting to different ecological niches in isolation. In time, these groups accumulated distinguishable morphological differences that some anthropologists have raised to the rank of different species [33], but speciation seems to be a lengthy process. Thus, molecular phylogenetic studies have found a long and consistent mean time to speciation in eukaryotes of around 2 million years (Myr) [34]. In fact, under climatic and demographic pressures these groups came into secondary contact several times. In some of these cases, recent ancient DNA (aDNA) studies have confirmed that, after separations of several hundred years, archaic groups as Neanderthals, Denisovans or Sima de los Huesos specimens hybridized frequently confirming the existence of incomplete sexual barriers among them[35–38]. The heads of these secondary encounters may be the exchange of genetic variation which greatly possibilities adaptation and avoids extinction. However, the tail of generalist species groups is that when coming into contact they have to compete for the same resources, so that the more adapted displace and outcompete the others with some genetic assimilation during this process [39]. Ultimately, the rate of assimilation or displacement depends on the amount of resources available. Thus, there is archaeological evidence that Neanderthals displaced less evolved erectus groups across Europe and archaic humans in the Middle East [40]; that in turn, modern humans displaced less adapted erectus groups in East and Southeast Asia [41], and Neanderthals in Europe [42,43]. Furthermore, again aDNA studies have demonstrated the extinction of several Neanderthal[44] and modern human populations [45,46] along its recent evolutionary history. From the above considerations I will consider all the groups described in this paper as sub-specific stages of a temporally evolving polytypic species.

### The northern African origin of the ancestor of modern humans and Neanderthals

Based on the topologies obtained from non-recombinant uniparental markers (Green et al. 2008; Mendez et al. 2016; Meyer et al. 2012, 2014; Petr et al. 2020)[35,36,47–49], I have proposed recently that modern humans and Neanderthals were sister clades [50], and that the topologies obtained using autosomal markers [37,48,51], which consider *Homo sapiens* as an outgroup of the sister pair Neanderthal-Denisovan were due to secondary introgression. Furthermore, I also posit that the ancestor of modern humans and Neanderthals originated in northern Africa, and that pre-Neanderthal groups crossed to Europe whereas the ancestors of modern humans remained in northern Africa, so that both groups evolved in allopatry [50].

### The northwest African origin of early anatomically modern humans

Human fossil remains and Middle Stone Age (MSA) archaeological artefacts from Jebel Irhoud, Morocco, dated at 315 ± 34 thousand years ago (Kya) have situated the earliest phase of modern human evolution in northwest Africa [4,5]. Applying a variable evolutionary rate dependent of temporal fluctuations in population size to the mtDNA genome, a coalescent age for the most recent common ancestor of all extant mtDNA lineages was estimated around 300 kya [14,21] which has been replicated in this study (Table 1).

**Table 1:**
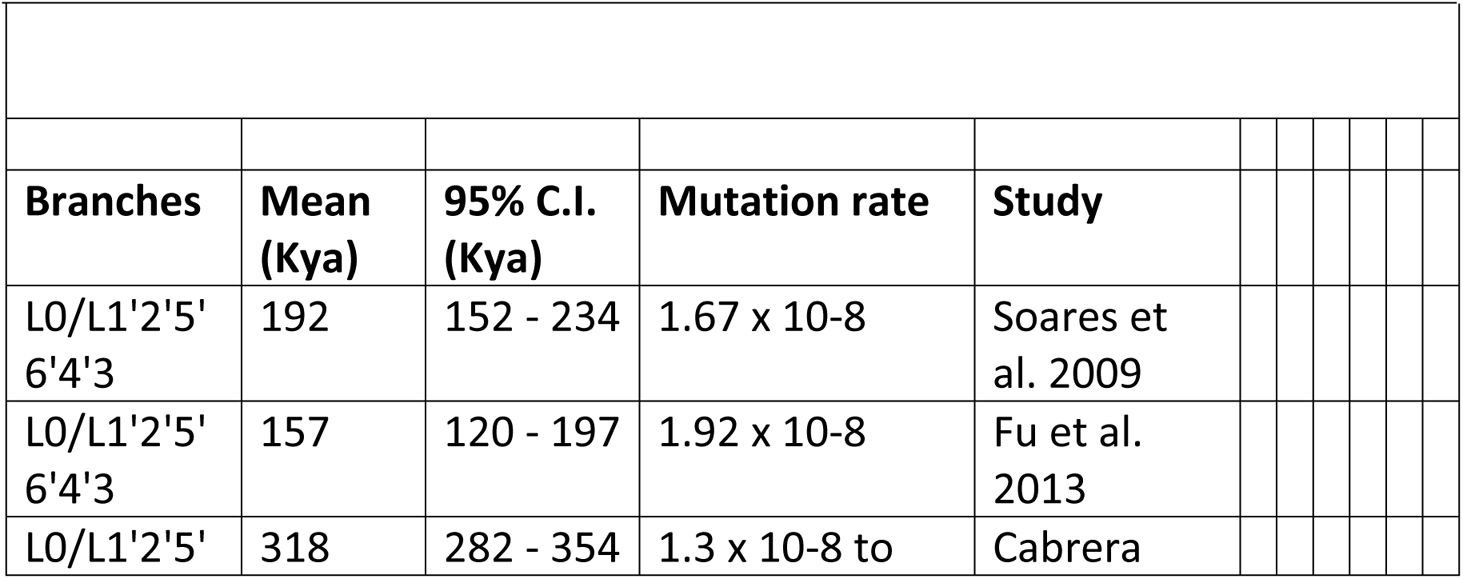
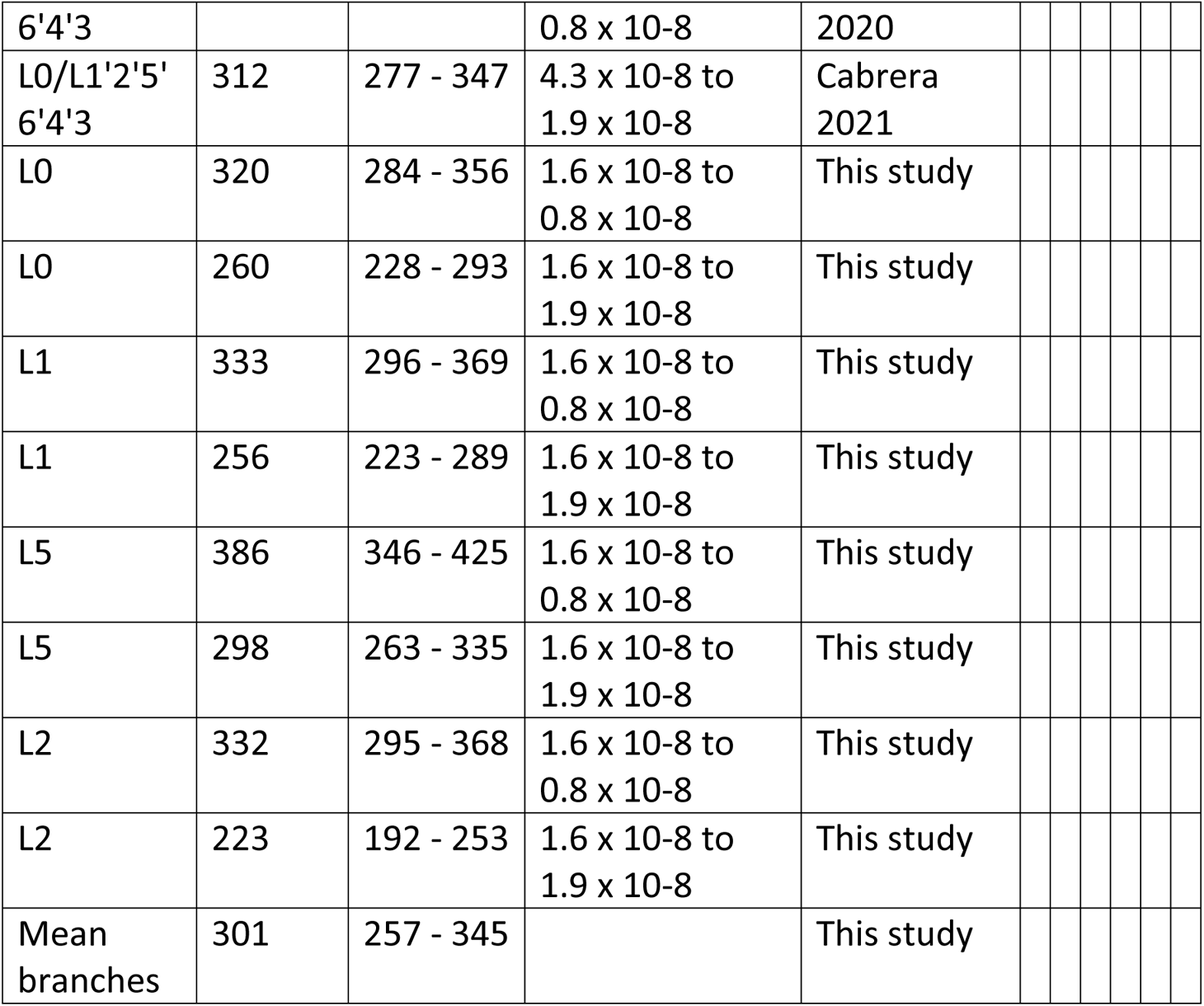
Mitochondrial DNA Estimate Ages to the most recent common ancestor of modern humans (L0/L1’2’5’6’4’3 split)

|  |  |  |  |  |  |  |  |  |  |
| --- | --- | --- | --- | --- | --- | --- | --- | --- | --- |
| Branches | Mean (Kya) | 95% C.I. (Kya) | Mutation rate | Study |  |  |  |  |  |
| L0/L1'2'5'6'4'3 | 192 | 152 - 234 | $1.67 \times 10^{-8}$ | Soares et al. 2009 | | | | | |
| L0/L1'2'5'6'4'3 | 157 | 120 - 197 | $1.92 \times 10^{-8}$ | Fu et al. 2013 | | | | | |
| L0/L1'2'5' | 318 | 282 - 354 | $1.3 \times 10^{-8}$ to | Cabrera | | | | | |

|  |  |  |  |  |  |  |  |  |  |  |
| --- | --- | --- | --- | --- | --- | --- | --- | --- | --- | --- |
| 6'4'3 | | | $0.8 \times 10^{-8}$ | 2020 | | | | | | |
| L0/L1'2'5'<br>6'4'3 | 312 | 277 - 347 | $4.3 \times 10^{-8}$ to<br>$1.9 \times 10^{-8}$ | Cabrera<br>2021 | | | | | | |
| L0 | 320 | 284 - 356 | $1.6 \times 10^{-8}$ to<br>$0.8 \times 10^{-8}$ | This study | | | | | | |
| L0 | 260 | 228 - 293 | $1.6 \times 10^{-8}$ to<br>$1.9 \times 10^{-8}$ | This study | | | | | | |
| L1 | 333 | 296 - 369 | $1.6 \times 10^{-8}$ to<br>$0.8 \times 10^{-8}$ | This study | | | | | | |
| L1 | 256 | 223 - 289 | $1.6 \times 10^{-8}$ to<br>$1.9 \times 10^{-8}$ | This study | | | | | | |
| L5 | 386 | 346 - 425 | $1.6 \times 10^{-8}$ to<br>$0.8 \times 10^{-8}$ | This study | | | | | | |
| L5 | 298 | 263 - 335 | $1.6 \times 10^{-8}$ to<br>$1.9 \times 10^{-8}$ | This study | | | | | | |
| L2 | 332 | 295 - 368 | $1.6 \times 10^{-8}$ to<br>$0.8 \times 10^{-8}$ | This study | | | | | | |
| L2 | 223 | 192 - 253 | $1.6 \times 10^{-8}$ to<br>$1.9 \times 10^{-8}$ | This study | | | | | | |
| Mean<br>branches | 301 | 257 - 345 |  | This study |  |  |  |  |  |  |

This mtDNA coalescence matches the archaeological and fossil estimates in Morocco but there is a lack of specific mtDNA lineages in this area to directly support a northwest African origin. However we have indirect evidence of the existence of an old genetic component in the Maghreb. Thus, Late Pleistocene northern African remains derived one-third of their genomic ancestry from a complex sub-Saharan African gene pool [52]. Curiously, this component was not detected in subsequent Neolithic periods [53]. On the other hand, it is interesting to point out that, although most of the Y-chromosome lineages in Morocco (J-M267; E-M81) are of recent implantation [54], one of the most ancient lineages of the Y-Chromosome, A0a1 (xP114) has been detected in Moroccan Berbers [55]. Accepting the northwest African origin hypothesis implies that other contemporaneous hominin lineages as the Broken Hill (Zambia) skull dated to 299 ± 25 kya [56], or the Kenyan Guomde calvarium dated to around 270 kya [57] possibly did not directly contribute to the origin of our species.

### The west central African southern African mtDNA bifurcation

The next phylogenetic step in the human mtDNA evolution was the split of the earliest L0 lineages from the L1’2’5’6’4’3 ancestor that seems to have occurred somewhere in central Africa around 230 kya (Table 2).

**Table 2.** Coalescent ages of the main African mtDNA haplogroups.

| Table 2. Coalescent ages of the main African mtDNA haplogroups |  |  |  |  |
| --- | --- | --- | --- | --- |
| Haplogroup | Lineages | Mutations | mean Age (years) | 95% C.I. (years) |
| L0 | 291 | 952 | 231,263 | 174,578 - 287,948 |
| L1 | 130 | 483 | 230,892 | 142,562 - 319,222 |
| L5 | 73 | 240 | 235,072 | 157,948 - 312,196 |
| L2 | 201 | 593 | 143,505 | 110,785 - 176,225 |
| L3'4'6 | 329 | 1,172 | 185,017 | 105,878 - 264,157 |
| L6 | 4 | 26 | 35,992 | 17,235 - 54,750 |
| L3'4 | 333 | 1,151 | 130,838 | 87,634 - 184,042 |
| L4 | 26 | 194 | 122,872 | 94,828 - 150,196 |
| L3 | 291 | 952 | 93,348 | 79,476 - 107,220 |

Subsequent subdivisions of L0 probably happened around the Zambezi river approximately 200 kya (Table 3)[58], whereas the L1 and L5 bifurcations occurred nearly at the same time in central Africa (Table 2).

**Table 3.**
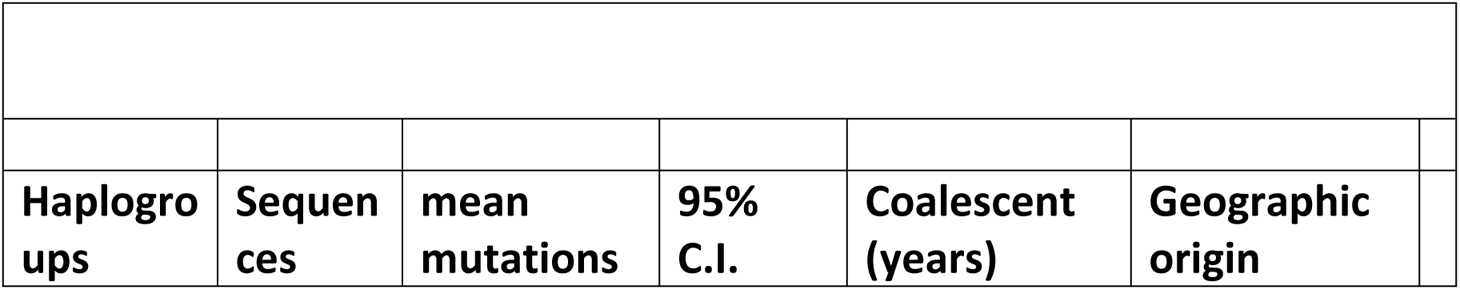
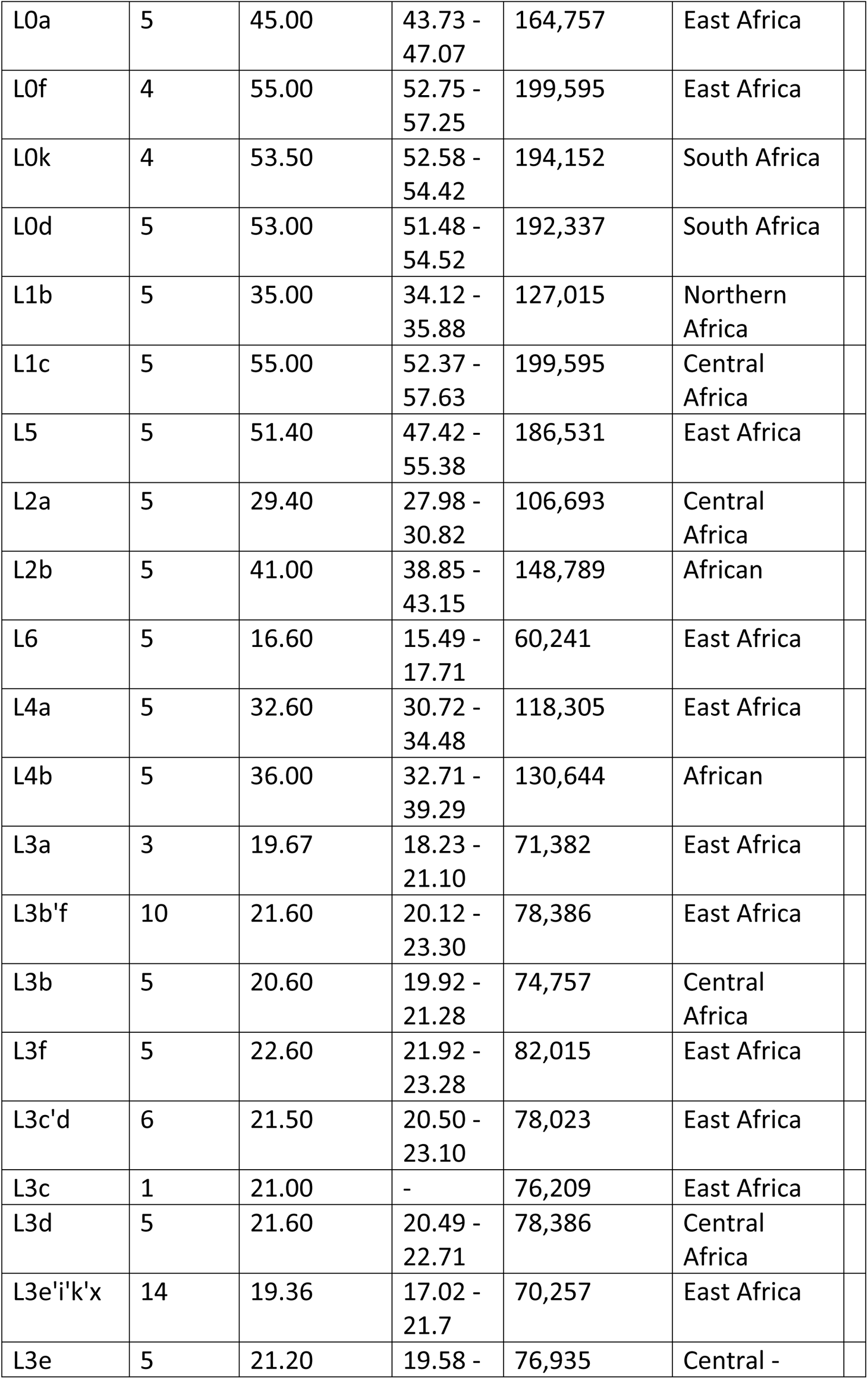
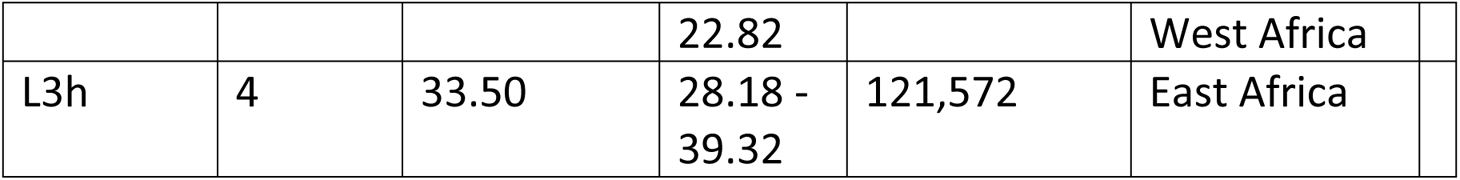
Mean number of mutations and probable geographic origin of the main African haplogroups.

|  |  |  |  |  |  |
| --- | --- | --- | --- | --- | --- |
| Haplogroups | Sequences | mean mutations | 95% C.I. | Coalescent (years) | Geographic origin |
| L0a | 5 | 45.00 | 43.73 -<br>47.07 | 164,757 | East Africa |
| L0f | 4 | 55.00 | 52.75 -<br>57.25 | 199,595 | East Africa |
| L0k | 4 | 53.50 | 52.58 -<br>54.42 | 194,152 | South Africa |
| L0d | 5 | 53.00 | 51.48 -<br>54.52 | 192,337 | South Africa |
| L1b | 5 | 35.00 | 34.12 -<br>35.88 | 127,015 | Northern<br>Africa |
| L1c | 5 | 55.00 | 52.37 -<br>57.63 | 199,595 | Central<br>Africa |
| L5 | 5 | 51.40 | 47.42 -<br>55.38 | 186,531 | East Africa |
| L2a | 5 | 29.40 | 27.98 -<br>30.82 | 106,693 | Central<br>Africa |
| L2b | 5 | 41.00 | 38.85 -<br>43.15 | 148,789 | African |
| L6 | 5 | 16.60 | 15.49 -<br>17.71 | 60,241 | East Africa |
| L4a | 5 | 32.60 | 30.72 -<br>34.48 | 118,305 | East Africa |
| L4b | 5 | 36.00 | 32.71 -<br>39.29 | 130,644 | African |
| L3a | 3 | 19.67 | 18.23 -<br>21.10 | 71,382 | East Africa |
| L3b'f | 10 | 21.60 | 20.12 -<br>23.30 | 78,386 | East Africa |
| L3b | 5 | 20.60 | 19.92 -<br>21.28 | 74,757 | Central<br>Africa |
| L3f | 5 | 22.60 | 21.92 -<br>23.28 | 82,015 | East Africa |
| L3c'd | 6 | 21.50 | 20.50 -<br>23.10 | 78,023 | East Africa |
| L3c | 1 | 21.00 | - | 76,209 | East Africa |
| L3d | 5 | 21.60 | 20.49 -<br>22.71 | 78,386 | Central<br>Africa |
| L3e'i'k'x | 14 | 19.36 | 17.02 -<br>21.7 | 70,257 | East Africa |
| L3e | 5 | 21.20 | 19.58 - | 76,935 | Central - |
|  |  |  | 22.82 |  | West Africa |
| L3h | 4 | 33.50 | 28.18 -<br>39.32 | 121,572 | East Africa |

Although L0, mainly the L0d’k clade, is considered a signature of the Khoe-San people [59], not all L0 branches remained in Southern Africa. Some of them as L0a and L0f had an early implantation in central and eastern Africa. Likewise, although L1c is representative of the western pygmy populations of central Africa, and L0a and L5 of the eastern ones [60], subsequent ramifications extended further to eastern Africa and beyond. Similarly, the two deepest Y-chromosome lineages, haplogroups A and B, branched out in these areas with some A lineages that seem autochthonous of central African pygmies as A1-P305, other of southern African Khoisan as A1b1a-M14, and other present in both areas as A1b1b-M32 [61–63]. The case of haplogroup B is similar, in fact Y-Hg B is a primary branch of the complex Y-Hg A [64], its deepest lineages in the B2-M182 clade are prevalent in western and central African pygmies, whereas other more derived branches as B2b1-P6 and B2b4-P8 are restricted to southern African Khoisan or to eastern Africans as B2b2 M169 [61–63]. Although subsequent expansions extended the geographic range of Y haplogroups A and B it seems that was in central Africa where both lineages originated [65].

The deep segregation of the ancestors of southern African populations from the rest was confirmed in a study of southern African ancient genomes in which the modern human divergence was estimated to 260 to 350 kya [66]. Furthermore, it was observed that differences between Khoisan genomes were greater than those between geographically very distant Eurasian genomes [67]. However, there is genomic evidence of secondary contacts among extant populations of Khoisan, rainforest pygmies, and click speakers Hadza and Sandawe from Tanzania which diverged by 100 – 120 kya [58,68]. In a similar vein, an ancient genomic divergence between the ancestors of the rainforest pygmies and West African Yoruba farmers was estimated to 90 – 150 Kya [69]. On the other hand, ancient DNA studies of a 2,330 years old South African skeleton observed the extinction of some L0d mtDNA lineages even in recent times confirming an evolution-extinction process in these populations [70].

In apparent contradiction with the mtDNA phylogeny, some genome based studies proposed that the deepest splitting branch in modern Africans leads to central African pygmies instead of southern African Khoisan [71], In my opinion, under a neutral view, this discrepancy can be explained as due to the non-recombining nature of the maternal lineages. Autosomal phylogenies are based on differences in genetic diversity, but in mtDNA, in addition, it is also based on the relative age of the mutations accumulated in the no recombining mtDNA genome. Due to this, although the central African pygmy L1c lineages show a mean number of mutations similar to, or greater than, the Khoisan L0 lineages (Table 3). The mtDNA phylogeny clearly shows that L0 is the deepest branch of the human tree (Sfig 1), and that the relative accumulation of mutations in the different lineages is, most probably, a result of their different demographic processes [14]. The surprising find that central African pygmies have reduced chromosome X to autosome diversity ratios relative to all other sub-Saharan Africans has also been explained by demography [12]. Thus, genetic age estimations situated the ancestor of modern humans in Central-Southern Africa in a temporal window contemporary with the Sangoan/Lupemban lithic technologies [72,73] and hominin specimens as Kabwe [56], Florisbad [74] or *Homo naledi* [75]. However, this phylogenetic jump from northwestern to central Africa leaves a geographic gap that is covered by western Africa. Regrettably, although its archaeological record is still scarce, western Africa seems to be a region of delayed and stagnant hominin evolution. Putative Oldowan, Acheulean, Sangoan and other Middle Stone Age incipient industries are sensibly more recent in West Africa[76–78] than their counterparts in eastern and central-southern Africa[7,79]. Likewise, the scarce hominin fossils remains unearthed show primitive features even at recent Pleistocene to Holocene boundaries as is the case of the Nigerian Iwo Eleru remains, reflecting either admixture with archaic humans or long-term survival of primitive anatomical features at recent (11.7-16.3 Ka) times[80,81]. On its hand, mtDNA does not present any deep split that could be specifically related to West Africa (Table 3). Nevertheless, the deepest Y-chromosome branch (Hg A00) with a coalescent age around 300 kya has been detected only in present day Cameroon populations, with particular prevalence among Mbo (6.3%) and Bangwa (40.3%) groups [82]. Furthermore, this basal lineage has been found in Late Pleistocene / Holocene forager remains from Shum Laka also in Cameroon [71]. It has been suggested that the presence of Hg A00 in modern humans could be the result of admixture with archaic hominins [71] but even if we excluded this haplogroup, there are other Y-chromosome basal lineages, as A0a1a observed in Cameroonian Bakola directly related to the A0a1 (xP114) present in Berbers from Algeria, or the primitive A1a clade observed in Fulbe and Tuareg from Niger and also found in Moroccan Berbers, that consistently points to an early migratory input from northwest to western Africa[64]. In addition, the genome-wide study of the above mentioned Shum Laka fossil specimens clearly showed that these individuals are most similar to the present-day Central African pygmies than to the actual Cameroonian populations. This fact is reinforced by the presence of the central African Y-chromosome B2b-M112 and mtDNA L1c haplotypes in those specimens [71]. These results could be explained as the result of a post-Pleistocene turnover of a primitive autochthonous West African population or, most probably, as the retraction and subsequent substitution of a previously much more large central African population as also could be the case for the southern Chad [83].

### The geographic northeast progression of the mtDNA phylogeny

Haplogroup L5 was the next branch splitting off from macro-haplogroup L5’2’6’4’3 at approximately 235 Kya (STable 4). However, its basal sub-branches (L5a, L5b, and L5c) suffered long periods of stagnation not having subsequent ramifications until favorable climatic conditions during the last interglacial period (130-74 Kya), around 30 Kya, and after the LGM, in Holocene times. The core geographic area for this haplogroup comprises Tanzania, Kenya, Ethiopia and southern Sudan [84,85]. However, secondary branches are predominant today in more specific regions or ethnic groups. For example, L5a1c1 is prevalent in Mbuti pygmies from central Africa, L5a1c2 concentrate in Kenya, and L5a2 and its subsequent radiations occur in southeastern African regions. Possible Y-chromosome counterparts of these early expansions through east and northward Africa could be the A3b2-M13 and B2a1-M218 lineages [86,87]. Even today, the geographic preeminence of L0, L1 and L5 basal lineages in southern, central and eastern Africa respectively, seems to be the remnants of a very ancient maternal structure in the African continent. It is interesting to compare this vision with the very similar results obtained from the analysis of ancient genome-wide genotype data from terminal Late Pleistocene and early Holocene African hunter gatherers that showed, in the same geographic area, a clinal pattern with individual genomes well represented by varying proportions of Central African pygmy, Southern African, and Ethiopian related ancestries [88]. However, in general, ancient DNA genome based studies focus on more recent population movements and turnovers and on the evidence of archaic introgression in the majority of the populations analyzed [71,89,90].

The wide chronological window open in eastern Africa by the L5 coalescent interval (95% CI: 312 – 158 kya) allows to include in it the most notable fossil and stone assemblages excavated in this area as are the modern human remains recovered at Omo-Kibish (Ethiopia) and dated to more than 200 kya[91,92], or the Herto (Ethiopia) remains dated around 160 kya[93]. The Sangoan-Lupemban lithic industries of equatorial Africa, mentioned above, have also been found at Lake Eyasi in Tanzania[94], and in Kenya at the Muguruk site[95], even most interesting is the presence of stratified Sangoan-Lupemban assemblages as far as northern Sudan, at Sai Island, dated around 230 kya that has been interpreted as the result of a possible human norward dispersal from equatorial Africa during the MIS 7 interglacial period[96,97].

### An earlier out of Africa

The next bifurcation in the mtDNA genomic tree separated two sister branches, L2 and the composite L3’4’6 with coalescent ages of 143 (111 – 176) kya and 185 (106 – 264) kya, respectively (Table 2). Based on its subsequent ramifications and present-day phylogeography, it has been suggested a western African origin for haplogroup L2 [98,99]. This seems to be in contradiction with the eastern geographic spread of its ancestral branch L5, and with the clear northeastern spread of its sister branch L3’4’6 [23,85,100]. However, as the earliest radiations of L2 occurred rather late, at around 60 kya, involving eastern and western expansions, it seems more equidistant to assume a central African origin for L2, an alternative hypothesis also contemplated by other authors[98,99]. In any case, haplogroup L2 is a typical sub-Saharan African lineage that likewise their predecessors L0, L1 and L5 did not participate in the out of Africa spread. It is the northeastern L3’4’6 cluster the progenitor of the entire Eurasian maternal diversity [23]. Haplogroup L6 was the first lineage to split off from that composite clade. This rare lineage presents mean frequencies below 1% in northeastern Africa but, in spite of this, it is found at similar frequencies in Saudi Arabia[23] and in higher frequencies in Yemen[101,102]. Based on the L6 tree[23] it appears that not all of the Arabian lineages are a subset of the African lineages, so that an early expansion of modern humans from Africa across Arabia has been suggested based on the haplogroup L6 phylogeography[101]. In a similar vein, haplogroup L4, another minor eastern African mitochondrial lineage, has Arabian representatives in all their main sub-branches, excepting L4b2b [23], which also points to an early phylogeographic extension of this clade into the Arabian Peninsula. The sister clade of L4 is haplogroup L3 that houses the Eurasian branches M and N which contain all of the mtDNA diversity outside Africa [103]. It has been proposed that after the radiation of L3 in eastern Africa, the ancestors of M and N crossed the Bab al Mandab strait about 60 – 70 Kya (the previously calculated coalescent age for L3) and, following a southern coastal route, they spread all over the world [100,104–106]. As an alternative hypothesis, we have proposed that the clade L3’4’6 already extended its geographic range to southwestern Asia and that the splits of the L6 and L3’4 branches (Table 2) occurred at the outside margins of Africa, being the Y-chromosome counterpart of this early spread the haplogroup CT-M168 that includes the Eurasian haplogroups C, D and F and the African haplogroup E[23]. The ample statistical range of these mtDNA coalescent ages (Table 2) includes important archaeological finds in the region as the presence of early modern human populations in the Levant at Misliya Cave from 177 to 194 kya[8], and at Qafzeh[107,108] and Skhul[109,110] caves from 90 to 130 kya. These findings are in support of an early expansion of modern humans from northeast Africa through the northern Levantine route[111] which has also been proposed by mtDNA [103,112] and genomic[113] data. However, fossil and previous genetic models propose different chronologies as the mtDNA and Y-chromosome dispersions are limited by the younger coalescent age of haplogroup L3 and CT-M168 respectively, and those based on genomic data by the levels of haplotype diversity of the population outside Africa and the genome mutation rate. In addition, it seems that the comparison of the lithic industries, prevalent in the areas implied in the two routes out of Africa, show stronger technological and typological similarities between assemblages from the Horn of Africa and the Nile Valley and Arabia than any of these regions and the Levant[114,115], however, alternatives to this vision exist [116]. On the other hand, mainly two MSA archaeological eastern African connections with Arabia have been identified, suggesting early expansions of modern humans from the former to the later. The first involves the Jebel Faya 1 site (United Arab Emirates) assemblage C, dated to about 125 kya[117], which lithic technologies show similarities with MSA assemblages in northeast Africa, particularly with the late Sangoan [118]. The second evidence is founded on the similarities of the Dhofar (Oman) lithic material and the Late Nubian Complex a specific African industry that in Dhofar is dated at 106 kya[119]. These potential arrivals coincide with wet stages of MIS5, with the split of the mtDNA L3’4 clade (Table 2), and also with genomic results that place indigenous Arabs as direct descendants of the first Eurasian populations [120], showing a comparative excess of Basal Eurasian ancestry [121]. However, recent archaeological sequences excavated in different regions of Arabia have evidenced hominin presence since 400 kya in the Nefud Desert [122], and since 210 kya at Jebel Faya[123] enabling much older hominin expansions into the Peninsula or even to an autochthonous hominin evolution in southwest Asia that got extinct by adverse climatic cycles and/or the arrival of modern humans. Finally, it should be mentioned that an exit through the Bab al Mandab Strait does not guarantee the existence of a southern coastal route since an inland northward expansion is also possible [124]. Furthermore, from the gathered evidence, both, the northern and southern migratory routes could have been followed alike [125]. At this respect, the detection of Nubian assemblages at the Negev highlands in the southern Levant dated to the MIS5 humid period is relevant [126].

### An earlier return to Africa

After a period of maturation and stasis in southwestern Asia, mtDNA haplogroup L3 split in the region and while the ancestors of the L3 African subclades returned to Africa, the ancestors of the Eurasian branches M and N began their exodus eastwards[23]. According to the new coalescent ages for the L3 subclades (Table 3), the first radiations in eastern Africa took place around 75 kya, at the beginning of the arid MIS 4 period. It was at this stage when an early modern human displacement by the Neanderthals in the Levant was attested [40]. The Y-chromosome counterpart of this mtDNA back flow to Africa was haplogroup E [127]. The detection, in the extant population of Saudi Arabia, of the basal African Y-chromosome lineage E-M96*[128] is in support of this back flow. Furthermore, whole genome sequence analyses also favor models involving possible African returns 70-60 kya[129,130]. Interestingly, a similar model, involving back flow to Africa, has been proposed to explain the complex mtDNA phylogeography of hamadryas baboons lineages present in Africa and Arabia [131].

The evidence gathered from the fossil and archaeological records for the proposed return to Africa has been only occasionally mentioned but, without generalized acceptance. Thus, it has been suggested that the Early Nubian Complex, developed at the end of MIS 6 beginning of MIS5 (145 – 125 Kya) in northeast Africa, extended to Arabia where the Late Nubian Complex occurred and from there was reintroduced into Africa during MIS5a (85-75 kya)[132]. It is known that an early Nubian technology appeared at Gademotta (Ethiopia) after 180 kya[133], and that it succeeds the Lupemban at Sai Island (Sudan) after 150 kya[96]. In addition, at Sodemein Cave (Egypt), stratigraphic layers dated to 121 ± 15 kya and 87 ± 9 kya are associated respectively with Early and Later Nubian complexes [134]. Potential modern human fossils coetaneous of these assemblages could be the Herto (Ethiopia) skull, dated to between 160 and 154 kya [135], and the Singa (Sudan) skull dated to 133 ± 2 kya[136]. Other Late Nubian assemblages could be mentioned, for instance, at Aduma (Ethiopia) where it is associated with skeletal remains dated to 79-105 kya [137] and in Taramsa Hill (Egypt) where it is at the same level of a child burial dated to 68.6 ± 8.0 kya[138]. After the dry MIS4, a transformed Nubian technology is present at Nazlet Khater (Egypt) associated with modern human skeletal remains dated to around 40 kya[139], already within a generalized MIS 3 population fragmentation that propitiated the cultural differentiation evidenced by the Later Stone Age African diversity. It should be emphasized that the proposed return to Africa, inferred from the non-recombinant maternal haplogroup L3 and paternal haplogroup E lineages, was earlier, had a broader geographic distribution, and greater genetic impact than later Eurasian penetrations into Africa. At this respect, it is suffice to note that, on average, maternal L3 lineages represent 27% and paternal E lineages 72% of the female and male African genetic pools respectively [23]. Subsequent pre-Holocene and Holocene Eurasian waves into Africa, signaled mainly by mtDNA haplogroups M1 and U6 [140–144], and Y-chromosome haplogroups J1-M267, R-V88 and T-M70 [87,145–147] had more limited impact affecting mainly northern and northeastern Africa. Due to the fact that these secondary Eurasian flows did not reach southern Africa, the delayed presence in South Africa of Nubian technology dated to 60-50 kya[148], and the analysis of the Hofmeyr skull, dated at 36.2 ± 3.3 kya, and showing strongest morphometric affinities with Upper Paleolithic Eurasians rather than present-day Khoisan [149], might be explained as the late arrival to the south of the proposed southwestern Asian reflux into Africa. The morphological affinities found between Hofmeyr and Nazlet Khater crania [150] are also in accordance with this hypothesis.

## Discussion

### Journey and evolution of modern humans throughout Africa

The proposition that the population from which modern humans evolved was located in northwest Africa is based on two main premises: first, it was the most probable place in which an ancestral hominin population bifurcated giving rise to the ancestors of the European Neanderthals and the African humans [50]; second, it has been there where the oldest remains of our species have been found [5].

Uniparental marker phylogenies point to Central/Southern Africa as the place where the first split of that population occurred. The association of these groups with Sangoan and Lupemban lithic technologies agree in time and space, however, it seems a cultural throwback that descendant of the makers of Mousterian MSA industries [4] opted for more primitive lithic strategies, although this could be justified as a special adaptation to new environments. At this respect, it should be mentioned the presence of a Sangoan of northeastern Africa technology included over a northwestern Africa Levallois Mousterian substratum at Wadi Lazalim in southern Tunisia [97]. Afterward, molecular markers signal a clear northward geographic progression signaled by L5 and L3’4’6 mtDNA clades at the eastern African region and, less evidently, by the L2 clade at the central region. In northeastern Africa it seems that the sub-Saharan Sangoan/Lupemban was replaced by the Early Nubian technology [132]. It is also probable that in northern Africa it was the Aterian which evolved from previous sub-Saharan lithic industries [151]. Nevertheless, the out of African migrants carrying maternal clade L3’4’6 and paternal clade CT-M168 only could brought an Early Levantine Mousterian industry to the Levant and a possible related Lupemban technology to southern Arabia [117] and, afterwards, an Early Nubian technology that spread and differentiated across the whole peninsula [152]. These early demic spreads out of Africa into Eurasia, coinciding with humid periods as the end of MIS7 (around 190 ka) and MIS5e (around 130 kya), could satisfactorily explain the detection of anatomically modern human teeth in southern China dated to 120-80 Kya[9], the presence of an early modern human tooth in Sumatra at 73-63 kya[153], the archaeological evidence of a possible human arrival to northern Australia around 65 kya [154], or the genomic evidence of an ancient split between Africans and Papuans around 120 kya[10].

Although the non-recombining uniparental markers have drawn a clear trajectory of modern humans across Africa, this certainly has not been the case. The presence of other primitive human groups along the way had undoubtedly promoted genetic admixture events that, unnoticed by uniparental markers, have been reflected in the genome of modern Africans [155,156] and their Eurasian descendants several times [157,158]. Furthermore, extinction events generated by simple genetic drift could affect more frequently to uniparental markers than whole genomes. Thus, some early demographic expansions detected by the analysis of complete genomes in current populations might not be perceived by the same analysis in uniparental markers. However, in spite these caveats, the phylogeny and phylogeography of mtDNA and Y-chromosome lineages seem to find a coherent reflection in the archaeological and anthropological records and might open the way for more detailed interdisciplinary studies.

A graphical map of the proposed modern human route and its cultural, physical, and genetic evolution across Africa is depicted in Fig 1.

**Fig 1.**
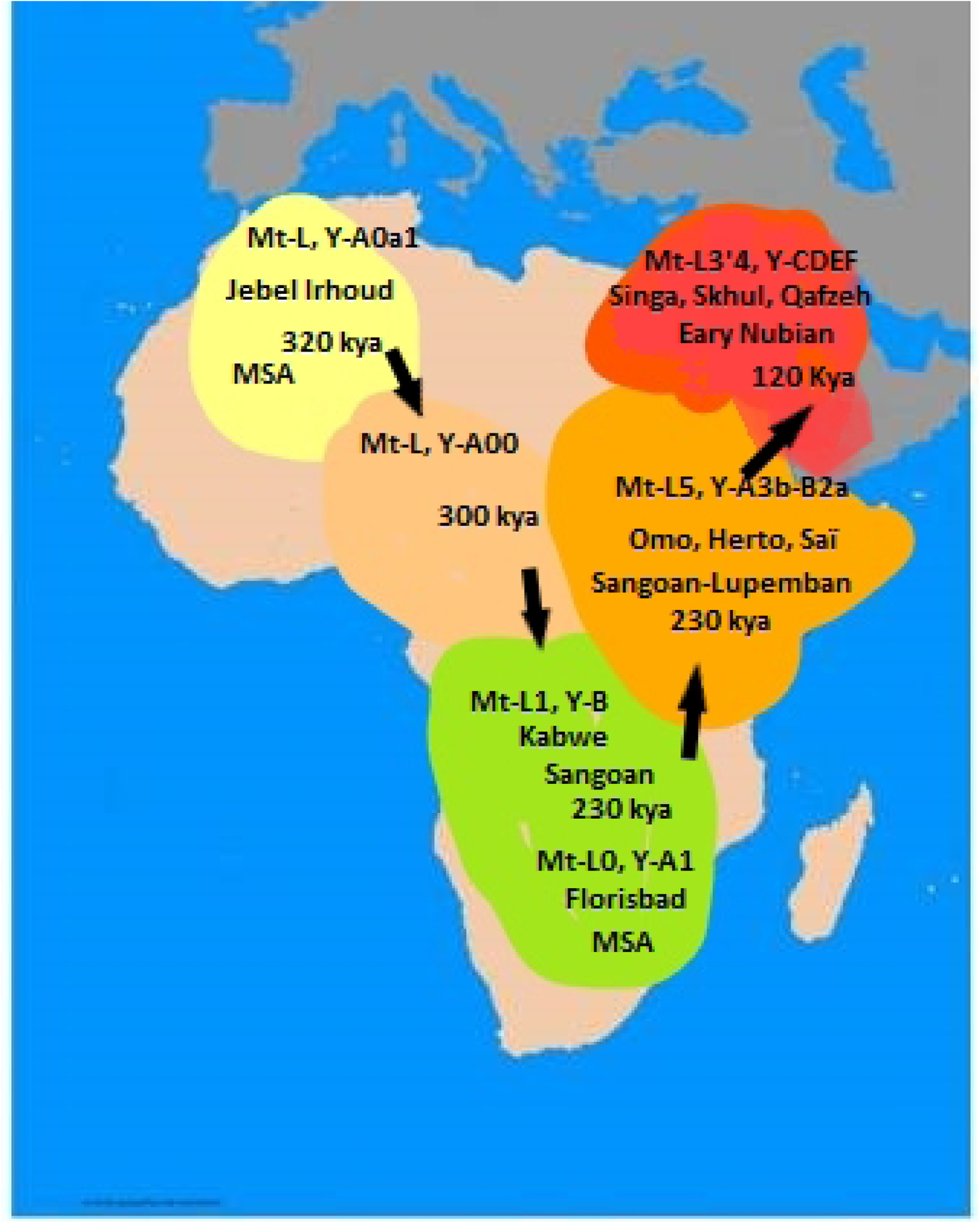
Modern human evolution across Africa and beyond.

### The first back to Africa of modern humans

The first out of Africa and back again for modern humans was proposed based on a nested cladistic analysis of the Y-chromosome variation (Hammer [159], and was supported by applying a most parsimonious criterion at an unbiased Y-chromosome tree [127]. Searching for a female counterpart, it was suggested that mtDNA haplogroup L3 also signals an early return to Africa [23] and, recently, this backflow to Africa has also been detected by whole genomic data [129]. The relatively closer morphological affinities of some African fossils with coetaneous Eurasian remains rather than with current African groups that have never abandoned the African continent [150], could also be taken in favor of this return to Africa. However, the archaeological support is much weaker because, although the temporal margins of the appearance and development of the Early and Late Nubian technological complexes are into the range proposed by the genetic markers, a clear geographical and temporal separation between these two lithic variants have not been yet determined. Therefore, it remains to deepen into this possibility suggested only by a few [132].

A graphical map of the proposed early return to Africa of modern humans and its genetic and archaeological support is depicted in Fig 2.

**Fig. 2.**
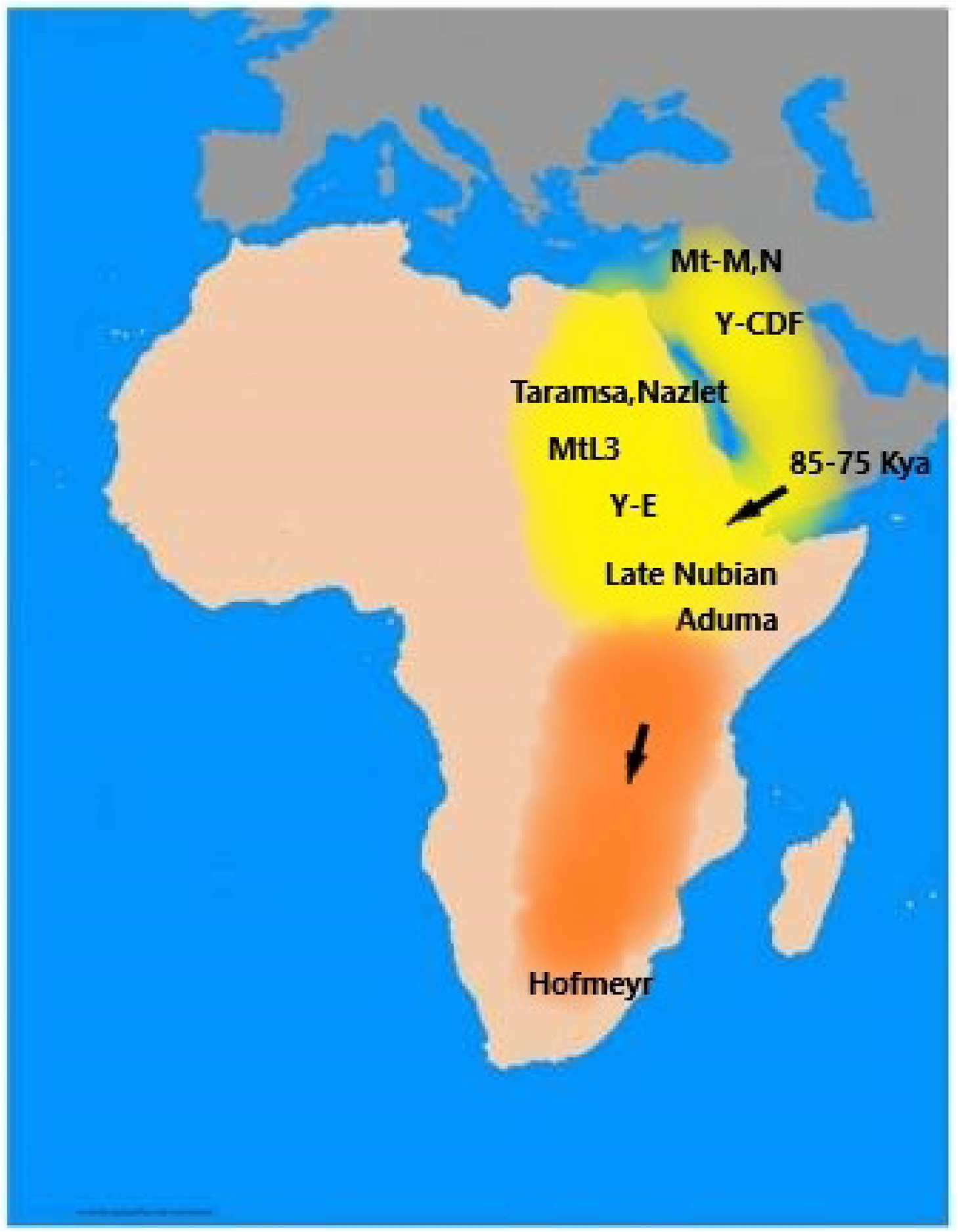
Modern human early return to Africa from southwest Asia.

## Supporting information captions

S1 Fig. Phylogenetic relationships between the main mtDNA African clades.

S2 Fig. Phylogeography of mtDNA haplogroup L2

S3 Fig. Phylogeography of mtDNA haplogroup L6

S4 Fig. Phylogeography of mtDNA haplogroup L4

S1 Table. Mitochondrial sequences utilized for the phylogenetic and phylogeographic analyses.

S2 Table. Age of the African mtDNA haplogroup L0.

S3 Table. Age of the African mtDNA haplogroup L1.

S4 Table. Age of the African mtDNA haplogroup L5.

S5 Table. Age of the African mtDNA haplogroup L2.

S6 Table. Age of the African mtDNA haplogroup L6.

S7 Table. Age of the African mtDNA haplogroup L4.

S8 Table. Age of the African mtDNA haplogroup L3.

